# Early Growth Assessment of *Treculia africana* Seedlings Influenced by Different Light Intensities and Mycorrhizal Inoculation

**DOI:** 10.1101/2022.10.04.510781

**Authors:** Oyefara Ayomide, Juliet Yisau

**Affiliations:** Federal University of Agriculture, Abeokuta, Nigeria, 111101

**Keywords:** Mycorrhiza, light intensity, *Treculia africana*, early growth performance

## Abstract

Light is essential for plant growth. The rate at which a plant grows depends on the light it receives. This study was carried out to observe the growth of *Treculia africana* seedlings under different light intensities (100%, 75%, 50%, and 25%) and mycorrhiza inoculation (Ectomycorrhiza, Endomycorrhiza, and topsoil {control}) with five replicates in each treatment. The experiment was laid out in a 3×4 factorial in CRD, and data were collected fortnightly for twelve (12) weeks. Data were subjected to analysis of variance in SAS, and the significant mean was separated using Fisher’s LSD. The result showed that the seedlings inoculated with ectomycorrhizal had the highest plant height (15.08), leaf area (26.90), number of leaves (5.75), collar diameter (2.61), fresh shoot weight (3.86), root weight (3.86), fresh weight (7.44), dry root weight (1.53), dry weight (2.87), absolute growth rate (0.34) and relative growth rate (0.39) which was significantly different from other treatment. Seedlings exposed to 75% and 25% light intensity had the highest performances compared to seedlings exposed to different light intensities. There was no significant difference (P>0.05) in the seedlings’ physiological variables assessed under the interaction of different light intensities and mycorrhiza inoculation. The seedlings inoculated with ectomycorrhizal and raised in 25% light intensity had the overall highest performance in all the growth variables assessed. Therefore, this study recommends raising *Treculia africana* seedlings with the inoculation of ectomycorrhizal and under shade to enhance early growth and massive nursery production.

## 1. Introduction

There are some nutrient-dense food crops are neglected in their cultivation but are high in protein, vitamins, and minerals. The crop species *Trecullia africana* is one of these. African breadfruit, or Treculia africana, is a multipurpose tree species. It grows in the forest zone in several tropical rainforests of West and Central Africa and is a member of the Moraceae family (Nzekwe & Amujiri, 2011). The enormous fruit and edible seeds of this plant, which can be boiled and consumed as starch like true breadfruit, are what give it the common name “African breadfruit.” In some rural communities of southeast Nigeria, it is a neglected and underutilized nutritional and medicinal crop with significant socioeconomic and cultural significance. The seeds are highly nutritious and constitute a cheap source of vitamins, minerals, proteins, carbohydrates, and fat (Osuji & Owei, 2010). Treculia africana is a large evergreen tree in tropical parts of Africa. It is one of the forest tree species known to be of immense domestic importance to both rural and urban dwellers (Enibe, 2007). The seeds are used for cooking; it also provides an excellent polyvalent dietetic value whose biological value exceeds that of soybean (WAC, 2004). The seeds are utilized in cooking, and they also have a superior polyvalent dietetic value that is superior to soybean in terms of biological value (WAC, 2004). The plant has various medicinal uses, including its cure for swellings, leprosy, cough, and rheumatism (WAC, 2005).

The mutual interactions between plants and microbes are essential in determining ecosystem productivity. The roots of most terrestrial plants form a symbiotic association with fungi known as MYCORRHIZAS. *Mycorrhiza* is a non-disease-producing association in which the fungus invades the root to absorb nutrients; mycorrhizal fungi establish a mild form of parasitism that is mutualistic, i.e. both the plant and the fungus benefit from the association. The fungus supplies nutrients, water, and minerals from the soil to the plant, and in return, the plant captures energy from the sun utilizing its chlorophyll and supplies it to the fungus. The fungi colonize the host plant root, form extension networks, and participate in the acquisition of phosphorus in the case of arbuscular mycorrhizal fungi (AMF) and nitrogen in ectomycorrhizal fungi (EMF) (Smith and Read 2008). Mycorrhizas increase root surface area for water and nutrient uptake. The use of mycorrhiza helps to improve higher branching of plant roots, and the mycorrhizal hyphae grow from the root to the soil enabling the plant roots to contact with the broader area of the soil surface, hence, increasing the absorbing area for water and nutrients absorption of the plant root system.

Light is one of the most crucial environmental factors required for plant growth. The intensity of light needed varies from species to species. Light is required for plant germination and growth. Alkaloids, glucosinolates, and cyanogen glycosides—metabolites that contain nitrogen—increase with lower light intensities (Coelho et al. 2007), whereas growth and plant biomass are best at higher light intensities (Wu et al. 2017). One of the most vital environmental elements for crop physiology and biochemistry is light quantity and quality (Yang et al., 2018a). For the majority of crop plants, even a small shift in light intensity causes significant changes in leaf morphology and structure (Wu et al. 2017).

*Treculia africana* is enlisted as an endangered species of Southern Nigeria and among the scantly researched indigenous fruit trees of African origin. Hence, this study examines the effect of light intensity and mycorrhizal inoculation on *Treculia africana*.

## 2. Materials and Methods

### Study Area

The study was carried out at the Forestry Nursery Unit of Forestry and Wildlife Management at the Federal University of Agriculture, Abeokuta, Ogun State, Nigeria. This area falls within latitudes 7010’N and 7058’N and longitudes 3020’E and 3037’E. It has a gently undulating landscape and a mild slope. The slope is punctuated by ridges, isolated, residual hills, valleys, and lowlands. The soils are sand and clay with a crystalline basement complex. It has an annual rainfall of 1200 mm with peaks in June and July; there is a dry season of two or three months. The area’s relative humidity is 82.54% and has an average monthly temperature of 35.80C. The forestry nursery is located about 50m from the campus.

### Experimental Design

The seeds of *Treculia africana were* sown on a germination bed and were watered daily for about three weeks. After germination, sixty seedlings were transplanted, and 20 of the seedlings were planted in polythene partly filled with topsoil, and 30 grams of arbuscular mycorrhiza were added and then filled with topsoil; another 20 of the seedlings were planted in a polythene pot filled with one head pan of Ectomycorrhiza thoroughly mixed with topsoil; and the last 20 seedlings were planted in a pot filled with topsoil, which served as the control. There were five replicates for each treatment, and a total of fifteen seedlings were placed in different light chambers.

Three light screening chambers were constructed for the experiments in such a way that wooden frames were built and then covered with layers of 1 mm mesh net on all sides except the side facing the ground; some researchers have demonstrated that a layer of 1 mm reduces the light intensity by 25% (Akinyele, 2007; Aderounmu, 2010; Akinyele & Dada, 2015). Therefore, treatment one was achieved by exposing the seedlings to direct sunlight (100% light intensity), which served as the control; treatment two was constructed with one layer of the mesh net (75% light intensity); treatment three was constructed with two layers of the mesh net (50% light intensity), and treatment four was constructed with three layers of the mesh net (25% light intensity).

The experiment was arranged in a 3 * 4 Complete Randomized Design. The following growth variables were assessed fortnightly for twelve weeks: stem height (cm), stem diameter (mm), leaf area (cm), and the number of leaves. A veneer calliper was used to measure the stem diameter; a measuring tape was used for the stem height, Leaf area = 4.41 + 1.14(L X B), while counting was used to determine the number of leaves. Physiological parameters were assessed at the end of the twelve weeks, and the following variables were assessed: fresh root weight, dry root weight, fresh shoot weight, dry shoot weight, total fresh weight, total dry weight, absolute growth rate, and relative growth rate.

### Data Analysis

Data were subjected to analysis of variance in SAS. The significant mean was separated using fisher’s LSD.

## 3. Results

### • Effect of mycorrhiza inoculation on morphological parameters of *Treculia africana* seedlings

Increments in seedlings’ morphological variables were highly affected by mycorrhiza inoculation. The number of leaves (5.75) was highly influenced in seedlings inoculated with Ectomycorrhiza, which was significantly different (*P<0.05*) from seedlings raised with topsoil, with the most negligible effect (4.56). Ectomycorrhiza is also significantly different (*P<0.05*) in the seedlings’ height (15.08) and leaf area (26.899) compared to seedlings raised in topsoil with a minor effect (10.51) and (21.721) for plant height and leaf area respectively. However, collar diameter had no significant difference (*P>0.05*) from seedlings raised in both Ectomycorrhiza and endomycorrhiza, but it is significantly different (*P<0.05*) from seedlings raised in topsoil (Table 1).

**Table i:** Effect of mycorrhiza inoculation on morphological parameters.

|  | Number of leaves | Plant Height<br>(cm) | Collar Diameter (cm) | Leaf Area<br>(cm<sup>2</sup>/plt) |
| --- | --- | --- | --- | --- |
| M<sub>1</sub> | 5.75<sup>a</sup> | 15.08<sup>a</sup> | 2.61<sup>a</sup> | 26.90<sup>a</sup> |
| M<sub>2</sub> | 4.81<sup>b</sup> | 11.95<sup>b</sup> | 2.44<sup>a</sup> | 22.54<sup>b</sup> |
| M<sub>3</sub> | 4.56<sup>b</sup> | 10.51<sup>b</sup> | 2.05<sup>b</sup> | 21.72<sup>b</sup> |

Means within a column with the same superscripts are not significantly different ( $p>0.05$ ), LSD $p=0.05$ M<sub>1</sub>- Ectomycorrhiza, M<sub>2</sub>- Endomycorrhiza, M<sub>3</sub>- Topsoil (control).

### • Effect of mycorrhiza inoculation on physiological parameters of *Treculia africana* seedlings

The seedlings have fresh shoot weight (3.86g), fresh root weight (3.86g), fresh weight (7.44g), dry shoot weight (1.34g), dry root weight (1.53g), dry weight (2.87g), root to shoot ratio (1.17g), absolute growth rate (0.34g) and relative growth rate (0.39g) were highest in soils inoculated with ectomycorrhizal. This effect was significantly different from seedlings inoculated with endomycorrhizal and topsoil with the least dry weight (1.38g), fresh shoot weight (2.33g), root weight (1.68g), fresh weight (4.01g), dry shoot weight (0.74g), dry root weight (0.63g), root to shoot ratio (0.82g), absolute growth rate (0.16g) and relative growth rate (0.29g). (Table 2).

**Table ii:**
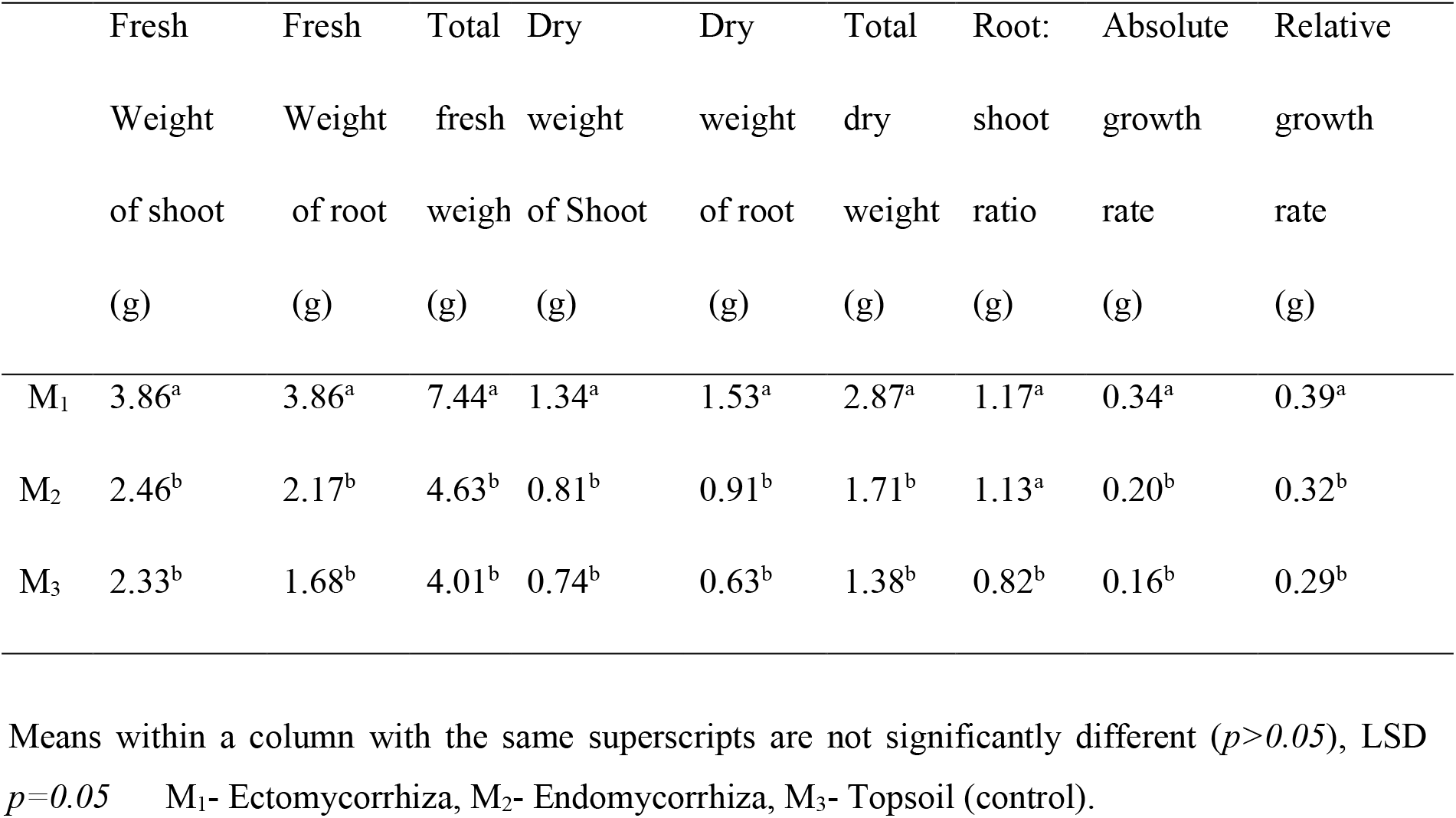
Effect of mycorrhiza inoculation on physiological parameters.

|  | Fresh Weight of shoot (g) | Fresh Weight of root (g) | Total fresh weigh (g) | Dry weight of Shoot (g) | Dry weight of root (g) | Total dry weight (g) | Root: shoot ratio (g) | Absolute growth rate (g) | Relative growth rate (g) |
| --- | --- | --- | --- | --- | --- | --- | --- | --- | --- |
| M<sub>1</sub> | 3.86<sup>a</sup> | 3.86<sup>a</sup> | 7.44<sup>a</sup> | 1.34<sup>a</sup> | 1.53<sup>a</sup> | 2.87<sup>a</sup> | 1.17<sup>a</sup> | 0.34<sup>a</sup> | 0.39<sup>a</sup> |
| M<sub>2</sub> | 2.46<sup>b</sup> | 2.17<sup>b</sup> | 4.63<sup>b</sup> | 0.81<sup>b</sup> | 0.91<sup>b</sup> | 1.71<sup>b</sup> | 1.13<sup>a</sup> | 0.20<sup>b</sup> | 0.32<sup>b</sup> |
| M<sub>3</sub> | 2.33<sup>b</sup> | 1.68<sup>b</sup> | 4.01<sup>b</sup> | 0.74<sup>b</sup> | 0.63<sup>b</sup> | 1.38<sup>b</sup> | 0.82<sup>b</sup> | 0.16<sup>b</sup> | 0.29<sup>b</sup> |

Means within a column with the same superscripts are not significantly different ( $p>0.05$ ), LSD $p=0.05$ M<sub>1</sub>- Ectomycorrhiza, M<sub>2</sub>- Endomycorrhiza, M<sub>3</sub>- Topsoil (control).

### • Effect of light intensity on morphological parameters of *Treculia africana* seedlings

The intensity of light greatly affected the morphological parameters of *Treculia africana* seedlings. Seedlings exposed to 25% light intensity (L4) had the highest number of leaves (5.42). This was significantly different (P0.05) in the collar diameter of the seedlings raised in 75% (l2), 50% (L3), and 25% (L4) light intensity compared to the seedlings raised in full sunlight 100% (L1) which had the least significant effect (2.11). However, the leaf area was not significantly different (P>0.05) across the different light intensities (Table 3).

**Table iii:** Effect of light intensity on morphological parameters.

|  | No. of<br>Leaf | Plant Height<br>(cm) | Collar<br>Diameter<br>(mm) | Leaf area<br>(cm<sup>2</sup>/plt) |
| --- | --- | --- | --- | --- |
| L<sub>1</sub> | 4.56<sup>b</sup> | 10.67<sup>b</sup> | 2.11<sup>b</sup> | 20.94<sup>a</sup> |
| L<sub>2</sub> | 5.22<sup>a</sup> | 15.23<sup>a</sup> | 2.50<sup>a</sup> | 25.62<sup>a</sup> |
| L<sub>3</sub> | 4.96<sup>ab</sup> | 11.93<sup>b</sup> | 2.37<sup>a</sup> | 24.31<sup>a</sup> |
| L<sub>4</sub> | 5.42<sup>a</sup> | 12.23<sup>b</sup> | 2.48<sup>a</sup> | 24.01<sup>a</sup> |

Means within a column with the same superscripts are not significantly different ( $p > 0.05$ ), LSD $p = 0.05$ L<sub>1</sub>- 100%, L<sub>2</sub>- 75%, L<sub>3</sub>- 50%, L<sub>4</sub>- 25%

### • Effect of light intensity on physiological parameters of *Treculia africana* seedlings

Seedlings of *Treculia africana* were highly affected by light intensity. Seedlings exposed to 25% light intensity (L4) had the highest fresh shoot weight (3.27g) and dry shoot weight (1.09g). These effects were not significantly different from seedlings raised under 100% light intensity (L1) (full sunlight) having the least effect on fresh shoot weight (2.51g) and the 50% light intensity (L3) had the least effect on dry shoot weight (0.85g). The seedlings exposed to 75% light intensity had the highest fresh root weight (2.80) which was not significantly different from the seedlings raised with 100%light intensity (L1) having the least effect (2.02g). Furthermore, seedlings exposed to 25% (L4) and 75 % (L2) light intensity had the highest fresh weight which is significantly different (P>0.05) from the seedlings exposed to 100% light intensity which had the least effect (4.52). However, dry root weight, total dry weight, root to shoot ratio, absolute growth rate, and relative growth rate was not significantly different (P>0.05) from the rate of light intensities. (Table 4).

**Table iv:** Effect of light intensity on physiological parameters.

|  | Fresh weight (%) | Fresh weight (%) | Total weight (%) | Dry weight (%) | Dry weight (%) | Total weight(g) | Root to shoot ratio(g) | Absolute growth rate (g/wk) | Relative growth rate (g/g/wk) |
| --- | --- | --- | --- | --- | --- | --- | --- | --- | --- |
| L<sub>1</sub> | 2.51<sup>a</sup> | 2.02<sup>a</sup> | 4.52<sup>b</sup> | 0.88<sup>a</sup> | 0.89<sup>a</sup> | 1.76<sup>a</sup> | 1.00<sup>a</sup> | 0.20<sup>a</sup> | 0.32<sup>a</sup> |
| L<sub>2</sub> | 3.04<sup>a</sup> | 2.80<sup>a</sup> | 5.84<sup>a</sup> | 1.05<sup>a</sup> | 1.06<sup>a</sup> | 2.11<sup>a</sup> | 1.02<sup>a</sup> | 0.25<sup>a</sup> | 0.34<sup>a</sup> |
| L<sub>3</sub> | 2.71<sup>a</sup> | 2.42<sup>a</sup> | 5.13<sup>ab</sup> | 0.85<sup>a</sup> | 0.98<sup>a</sup> | 1.83<sup>a</sup> | 1.08<sup>a</sup> | 0.21<sup>a</sup> | 0.32<sup>a</sup> |
| L<sub>4</sub> | 3.27<sup>a</sup> | 2.67<sup>a</sup> | 5.84<sup>a</sup> | 1.09<sup>a</sup> | 1.15<sup>a</sup> | 2.23<sup>a</sup> | 1.06<sup>a</sup> | 0.26<sup>a</sup> | 0.35<sup>a</sup> |

Means within a column with the same superscripts are not significantly different ( $p>0.05$ ), LSD $p=0.05$ L<sub>1</sub>- 100%, L<sub>2</sub>- 75%, L<sub>3</sub>- 50%, L<sub>4</sub>- 25%

### • Interactive effect of light intensity and mycorrhiza inoculation on morphological parameters of *Treculia africana* seedlings

The combined effect of light intensity and mycorrhiza inoculation enhanced morphological parameters of *Treculia africana* seedlings. The interactive effect of light intensity and mycorrhiza shows that the seedlings inoculated with ectomycorrhizal under 25% light intensity (l4) had the highest number of leaves (6.77) while the least number of leaves (4.03) was recorded in seedlings inoculated with topsoil with 50% light intensity. There is no significant difference in seedlings inoculated with mycorrhiza in 100% light intensity (L1). Furthermore, there is no significant difference (P>0.05) in the interaction of light and mycorrhiza on the seedlings’ height. The highest collar diameter (2.99) was recorded in 25% light intensity inoculated with ectomycorrhizal with the least collar diameter (1.76) recorded in 50% light intensity inoculated with topsoil. However, there is no significant difference in seedlings raised with ectomycorrhizal under different light intensities except for 100% light intensity. Lastly, the highest leaf area (32.10) was recorded in 25% light intensity inoculated with ectomycorrhizal and the least leaf area (17.58) was recorded in 50% light intensity inoculated with topsoil (Table 5).

**Table v:** Interactive effect of light intensity and mycorrhiza inoculation on morphological parameters.

| Light Intensity | Mycorrhiza inocula | Number of leaves | Plant Height<br>(cm) | Collar Diameter (mm) | Leaf<br>area (cm<sup>2</sup>/plt) |
| --- | --- | --- | --- | --- | --- |
| L<sub>1</sub> | M<sub>1</sub> | 4.77<sup>b</sup> | 11.81<sup>a</sup> | 2.13<sup>a</sup> | 20.92<sup>c</sup> |
|  | M<sub>2</sub> | 4.33<sup>b</sup> | 10.74<sup>a</sup> | 2.32<sup>a</sup> | 20.76<sup>c</sup> |
|  | M<sub>3</sub> | 4.57<sup>b</sup> | 9.47<sup>a</sup> | 1.88<sup>a</sup> | 21.13<sup>c</sup> |
| L<sub>2</sub> | M<sub>1</sub> | 5.70<sup>a</sup> | 18.74<sup>a</sup> | 2.52<sup>a</sup> | 25.00<sup>b</sup> |
|  | M<sub>2</sub> | 4.97<sup>b</sup> | 13.17<sup>a</sup> | 2.53<sup>a</sup> | 24.04<sup>b</sup> |
|  | M<sub>3</sub> | 5.00<sup>ab</sup> | 13.78<sup>a</sup> | 2.43<sup>a</sup> | 27.82<sup>a</sup> |
| L<sub>3</sub> | M<sub>1</sub> | 5.77<sup>a</sup> | 14.79<sup>a</sup> | 2.79<sup>a</sup> | 29.57<sup>a</sup> |
|  | M<sub>2</sub> | 5.07<sup>ab</sup> | 12.79<sup>a</sup> | 2.57<sup>a</sup> | 25.79<sup>b</sup> |
|  | M<sub>3</sub> | 4.03<sup>b</sup> | 8.20<sup>a</sup> | 1.76<sup>a</sup> | 17.58<sup>c</sup> |
| L<sub>4</sub> | M<sub>1</sub> | 6.77<sup>a</sup> | 14.97<sup>a</sup> | 2.99<sup>a</sup> | 32.10<sup>a</sup> |
|  | M<sub>2</sub> | 4.87<sup>b</sup> | 11.11<sup>a</sup> | 2.33<sup>a</sup> | 19.57<sup>c</sup> |
|  | M<sub>3</sub> | 4.63<sup>b</sup> | 10.60<sup>a</sup> | 2.13<sup>a</sup> | 20.36<sup>c</sup> |

Means within a column with the same superscripts are not significantly different ( $p>0.05$ ), LSD $p=0.05$ . M<sub>1</sub>- Ectomycorrhiza, M<sub>2</sub>- Endomycorrhiza, M<sub>3</sub>- Top soil (control). L<sub>1</sub>- 100%, L<sub>2</sub>- 75%, L<sub>3</sub>- 50%, L<sub>4</sub>- 25%

### • The interaction of light and mycorrhiza had no significant effect on the physiological parameters of *Treculia africana seedlings*

The seedlings inoculated with ectomycorrhizal and under 25% light intensity (L4) had the highest shoot weight (4.59), fresh weight (8.49), dry shoot weight (1.62), dry root weight (1.75), total dry weight (3.38), absolute growth rate (0.41) and relative growth rate (0.41). The seedlings inoculated with ectomycorrhizal under 75% light intensity had the highest root weight (4.05g). The highest root-to-shoot weight was observed in the seedlings inoculated with endomycorrhizal under 50% light intensity (L3). (Table 6).

**Table vi:** Interactive effect of light intensity and mycorrhiza inoculation on physiological parameters.

| LIGHT | MYCO | SW (g) | RW (g) | FW (g) | DSW | DRW | TDW | R: S | AGR | RGR |
| --- | --- | --- | --- | --- | --- | --- | --- | --- | --- | --- |
|  |  |  |  |  | (g) | (g) | (g) | (g) | (g/wk) | (g/g/wk) |
| L<sub>1</sub> | M<sub>1</sub> | 3.21<sup>a</sup> | 2.92<sup>a</sup> | 6.13<sup>a</sup> | 1.43<sup>a</sup> | 1.22<sup>a</sup> | 2.34<sup>a</sup> | 1.09<sup>a</sup> | 0.28<sup>a</sup> | 0.36<sup>a</sup> |
|  | M<sub>2</sub> | 2.23<sup>a</sup> | 1.99<sup>a</sup> | 4.21<sup>a</sup> | 0.80<sup>a</sup> | 0.93<sup>a</sup> | 1.73<sup>a</sup> | 1.22<sup>a</sup> | 0.20<sup>a</sup> | 0.32<sup>a</sup> |
|  | M<sub>3</sub> | 2.09<sup>a</sup> | 1.17<sup>a</sup> | 3.26<sup>a</sup> | 0.71<sup>a</sup> | 0.52<sup>a</sup> | 1.22<sup>a</sup> | 0.69<sup>a</sup> | 0.14<sup>a</sup> | 0.28<sup>a</sup> |
| L<sub>2</sub> | M<sub>1</sub> | 3.99<sup>a</sup> | 4.05<sup>a</sup> | 8.04<sup>a</sup> | 1.46<sup>a</sup> | 1.63<sup>a</sup> | 3.09<sup>a</sup> | 1.21<sup>a</sup> | 0.37<sup>a</sup> | 0.40<sup>a</sup> |
|  | M<sub>2</sub> | 2.57<sup>a</sup> | 1.72<sup>a</sup> | 4.29<sup>a</sup> | 0.80<sup>a</sup> | 0.65<sup>a</sup> | 1.46<sup>a</sup> | 0.82<sup>a</sup> | 0.17<sup>a</sup> | 0.31<sup>a</sup> |
|  | M<sub>3</sub> | 2.57<sup>a</sup> | 2.62<sup>a</sup> | 5.18<sup>a</sup> | 0.87<sup>a</sup> | 0.89<sup>a</sup> | 1.76<sup>a</sup> | 1.03<sup>a</sup> | 0.20<sup>a</sup> | 0.33<sup>a</sup> |
| L<sub>3</sub> | M<sub>1</sub> | 3.66<sup>a</sup> | 3.46<sup>a</sup> | 7.12<sup>a</sup> | 1.15<sup>a</sup> | 1.50<sup>a</sup> | 2.65<sup>a</sup> | 1.25<sup>a</sup> | 0.32<sup>a</sup> | 0.37<sup>a</sup> |
|  | M<sub>2</sub> | 2.54<sup>a</sup> | 2.68<sup>a</sup> | 5.22<sup>a</sup> | 0.86<sup>a</sup> | 1.07<sup>a</sup> | 1.93<sup>a</sup> | 1.27<sup>a</sup> | 0.22<sup>a</sup> | 0.34<sup>a</sup> |
|  | M<sub>3</sub> | 1.94<sup>a</sup> | 1.12<sup>a</sup> | 3.06<sup>a</sup> | 0.54<sup>a</sup> | 0.38<sup>a</sup> | 0.91<sup>a</sup> | 0.71<sup>a</sup> | 0.10<sup>a</sup> | 0.24<sup>a</sup> |
| L<sub>4</sub> | M<sub>1</sub> | 4.59<sup>a</sup> | 3.89<sup>a</sup> | 8.49<sup>a</sup> | 1.62<sup>a</sup> | 1.75<sup>a</sup> | 3.38<sup>a</sup> | 1.13<sup>a</sup> | 0.41<sup>a</sup> | 0.41<sup>a</sup> |
|  | M<sub>2</sub> | 2.50<sup>a</sup> | 2.30<sup>a</sup> | 4.80<sup>a</sup> | 0.77<sup>a</sup> | 0.95<sup>a</sup> | 1.72<sup>a</sup> | 1.21<sup>a</sup> | 0.20<sup>a</sup> | 0.33<sup>a</sup> |
|  | M<sub>3</sub> | 2.73<sup>a</sup> | 1.80<sup>a</sup> | 4.54<sup>a</sup> | 0.86<sup>a</sup> | 0.74<sup>a</sup> | 1.60<sup>a</sup> | 0.84<sup>a</sup> | 0.19<sup>a</sup> | 0.31<sup>a</sup> |

Means within a column with the same superscripts are not significantly different ( $p>0.05$ ), LSD $p=0.05$ . M<sub>1</sub>- Ectomycorrhiza, M<sub>2</sub>- Endomycorrhiza, M<sub>3</sub>- Topsoil (control). L<sub>1</sub>- 100%, L<sub>2</sub>- 75%, L<sub>3</sub>- 50%, L<sub>4</sub>- 25%
FW- Fresh weight, SW- Shoot weight, RW- Root weight, DSW- dry shoot weight, DRW- dry root weight, TDW- total dry weight, R TO S- root to shoot ratio, AGR- absolute growth rate, RGR- relative growth rate

## 4. Discussion

### † Effect of mycorrhiza inoculation on morphological and physiological parameters of *Treculia africana* seedlings

This study showed that mycorrhiza is an important treatment that influences the seedling growth of *Treculia africana*. This effect was observed in the number of leaves produced, increased stem diameter, plant height, and leaf area. This observation might be due to the presence of the mycorrhiza symbiotic inoculation, which enhances the growth of a plant. This was similar to the findings of Oyun *et al*. (2010), who reported that mycorrhiza inoculation enhanced the growth of *Acacia Senegal* both in the nursery and field. Also, Mi-Yeong and Ahn-Heum (2006) observed that increasing the number of ectomycorrhizal fungi would increase the seedling growth of *Pinus densiflora*. According to Ezenwenyi *et al*. (2020), the presence of mycorrhiza in soil influenced the growth performance of *Treculia africana*.

*Treculia africana* seedlings inoculated with mycorrhiza were more highly influenced than seedlings without mycorrhiza inoculation. The seedlings’ fresh weight, dry weight, shoot weight, dry shoot weight, root weight, dry root weight, relative growth rate, root to shoot weight, and absolute growth rate were more highly influenced by the inoculation of ectomycorrhizal than seedlings not inoculated with mycorrhiza. This could be associated with the higher accumulation of phosphorus, nitrogen, and potassium from the soil (Erman*et al*. 2011).

### † Effect of light intensity on morphological and physiological parameters of *Treculia africana* seedlings

Light affects several plant metabolic activities, such as photosynthesis and respiration. It is, therefore, essential to establish seedlings within the range of their tolerance level (Hongfang Zhu *et al.*). The study observed the influence of light intensity on the growth of *Treculia africana* seedlings. Seedlings’ height, collar diameter, and leaf area had the highest growth rates at 75% of the intensity of light. According to Adesokan *et al*. (2020), *Acacia muricata* requires partial shade (75% light intensity) for seedling growth and development in proper plantation establishment. Based on the study, *Treculia africana* requires some shade for the establishment and early growth, which agrees with what was reported on the light intensity requirement of *Chrysophylum albidum* (Onyekwelu *et al*. 2013). However, it was observed that 25% light intensity affected the shoot weight, dry shoot weight, dry root weight, dry weight, absolute growth rate, and relative growth rate. This observation was supported by Mattana *et al*. (2006), who reported higher biomass with reduced light intensity. It, however, contracts the findings of Mukhtar (2016), who reported that *Balanites aegyptiaca* had the highest dry weight when exposed to full sunlight.

### † Interactive effect of light intensity and mycorrhiza inoculation on morphological and physiological parameters of *Treculia africana* seedlings

The combined effect of light intensity and mycorrhiza inoculation had no significant influence on the seedlings’ morphological variables except the number of leaves produced. Seedlings inoculated with ectomycorrhizal under 25% intensity of light produced more leaves. This finding is supported by the observations of Miller (2014), who, in his study, observed that Lima Bean seedlings inoculated with mycorrhiza under full sunlight had less performance than seedlings under low light intensity inoculated with mycorrhiza. The study showed that the combined effect of light and mycorrhiza inoculation did not significantly influence seedling physiological parameters. On the contrary, Ballhorn *et al*. (2016) observed that plant development and reproduction are reduced in inoculated plants under shaded conditions and reproduction in inoculated plants under reduced light.

## 5. Conclusion

Mycorrhiza inoculation and light intensity play a vital role in the performance and early growth of *Treculia africana* seedlings. The study clarifies that early growth of *Treculia africana* was increased with the inoculation of Ectomycorrhizal and light intensities of 75% and 25%. However, it was discovered that seedlings inoculated with ectomycorrhizal and raised under 25% light intensity produced more vigorous seedlings than other treatments. In sum, it is observed from this study that *Treculia africana* requires shade for seedling growth and development. Also, mycorrhiza inoculation, especially ectomycorrhiza, brings about better growth performance of *Treculia africana*.

## 6. Authors Contribution

Oyefara Ayomide: carried out the research experiment

Dr. Juliet Yisau: contributed in terms of expertise and guidance

